# Rank concordance of polygenic indices: Implications for personalised intervention and gene-environment interplay

**DOI:** 10.1101/2022.05.03.490435

**Authors:** Dilnoza Muslimova, Rita Dias Pereira, Stephanie von Hinke, Hans van Kippersluis, Cornelius A. Rietveld, S. Fleur W. Meddens

## Abstract

Polygenic indices (PGIs) are increasingly used to identify individuals at high risk of developing diseases and disorders and are advocated as a screening tool for personalised intervention in medicine and education. The performance of PGIs is typically assessed in terms of the amount of phenotypic variance they explain in independent prediction samples. However, the correct *ranking* of individuals in the PGI distribution is a more important performance metric when identifying individuals at high genetic risk. We empirically assess the rank concordance between PGIs that are created with different construction methods and discovery samples, focusing on cardiovascular disease (CVD) and educational attainment (EA). We find that the rank correlations between the constructed PGIs vary strongly (Spearman correlations between 0.17 and 0.94 for CVD, and between 0.40 and 0.85 for EA), indicating highly unstable rankings across different PGIs for the same trait. Simulations show that measurement error in PGIs is responsible for a substantial part of PGI rank discordance. Potential consequences for personalised medicine in CVD and research on gene-environment (G×E) interplay are illustrated using data from the UK Biobank.

## Introduction

Since the publication of the first genome-wide association study (GWAS) in 2005, it has become clear that most common human behavioural and disease traits are polygenic: they are influenced by thousands of single nucleotide polymorphisms (SNPs), each with a tiny effect^1,2^. GWAS estimates can be used to calculate an individual’s genetic risk or predisposition using a polygenic index (PGI; also known as a “polygenic (risk) score”): a weighted sum of SNPs, with the weights proportional to the effect size estimates obtained from a GWAS in an independent sample^3,4^. The recent increase in the predictive power of PGIs has opened the door to their usage in clinical settings^5–7^. For example, one study found that individuals ranking in the top quintile of the PGI distribution for cardiovascular disease are most likely to benefit from statin treatment, lowering the 10-year relative risk of coronary heart disease by 45%, and no risk reduction for individuals in the lowest PGI quintile^6^. More controversially, PGIs are also starting to be used for embryo selection^8–10^, and it has been suggested that in the future PGIs might be used to select against embryos predisposed to learning disorders^11,12^.

While the performance of a PGI is typically assessed by its explained phenotypic variance in an independent prediction sample^11^, a PGI’s precision in correctly *ranking* individuals in the PGI distribution is arguably more important when using PGIs for personalised interventions. In personalised interventions, individuals at elevated genetic risk are typically identified by their rank in the PGI distribution (e.g., top quintile). Moreover, ranking precision is also likely to be important for the estimation of gene-by-environment (G×E) interplay. G×E studies analyse heterogeneity in treatment effects as a function of individuals’ PGI: this is possible using the full (continuous) PGI distribution or on the basis of quantile-stratified samples of the PGI distribution (e.g., above/below the median)^13–16^. Imprecise PGI rankings may therefore lead to noisy decision-making in the clinic and bias our understanding of G×E interplay.

While recent studies have started to stress the importance of transparency about the construction of PGIs^17–20^, empirical studies often implicitly assume that PGIs for a specific trait are interchangeable. PGIs can be constructed in various ways. The most important features that have been highlighted are (i) the choice of GWAS discovery sample, (ii) the number of SNPs included, and (iii) the weights used to construct the aggregated polygenic index – e.g., corrected for linkage disequilibrium or not^21^. In this study, we empirically analyse individuals’ rank concordance across PGIs with different construction methods and discovery samples and explore the mechanisms underlying the discordance. Rank discordance between PGIs could arise from differences between construction methods, differences in the environmental context of the discovery samples, or random measurement error stemming from the finite discovery samples. In the empirical analyses, we focus on the first two of these: the discovery sample and the construction method. We use simulations to explore the extent to which measurement error in the PGIs is responsible for PGI rank discordance.

We start by investigating individuals’ rank concordance for two different polygenic traits that have been highlighted as promising targets for personalised screening: cardiovascular disease (CVD)^5,6,22,23^ and educational attainment (EA)^11,12^. We compare the PGIs constructed using different discovery samples (i.e., UK Biobank (UKB)^24^, CARDIoGRAM^25^, and 23andMe, Inc.^26^) as well as the two most commonly used construction methods: the “clumping and thresholding” algorithm as implemented in Plink (henceforth C+T)^27,28^ and the Bayesian LDpred method^29^. C+T is widely used given its relative simplicity and low computational cost^21^, although Bayesian methods such as LDpred are gaining popularity due to the increased predictive power compared to C+T. The central difference between Plink C+T and LDpred lies in the fact LDpred corrects the SNP weights for linkage disequilibrium (LD) and uses a large set of SNPs (i.e., >1 million) in the PGI, whereas Plink deals with LD by only keeping one SNP from each LD block — typically the SNP with the lowest *p* value. Because the differences between different Bayesian PGI construction methods like LDpred, PRS-CS, or S-Bayes-R are relatively less pronounced^30^, they are not explicitly considered in this paper.

We find limited concordance in individuals’ ranking across PGIs that are created with different construction methods or discovery samples. For example, for EA, 17% of the individuals who are in the top quintile of the UKB-based PGI (C+T) are in the bottom quintile of the 23andMe-based PGI (C+T). For LDpred-based PGIs constructed on the basis of the same discovery samples, the switch from top to bottom quintile is around 9%. We present two applications to illustrate the impact of such rank discordance. First, we show how PGI rank discordance can affect treatment decisions. Following an earlier study^22^, we assess which individuals should be given statin treatment by combining classical CVD risk factors and CVD PGIs in a prediction model. Here, we find that treatment decisions can vary highly depending on the PGI used in the model. Second, inspired by a study^31^ analysing how genetic effects on EA (among other traits) differ across birth cohorts, we investigate G×E interactions using different EA PGIs and birth year. We show that choices in the PGI construction stage can affect G×E estimations and with that, our understanding of the interplay between nature and nurture.

To understand the source of PGI rank discordance, we conduct simulations that show how accurately one can identify an individual’s true PGI rank and phenotype rank under various degrees of measurement error in the PGI. Our simulations indicate that for a trait with a SNP-heritability of 25% and a PGI that explains 12% of the total phenotypic variation (i.e., the ‘explained SNP-heritability’ is about 50%; this is roughly the state-of-the-literature for EA^26^), we only classify half of the individuals correctly in the top PGI quintile. Importantly, we show that the rank misclassification depends on the *explained* SNP-based heritability and is largely independent of the *absolute* SNP-based heritability. Overall, the simulations show that measurement error in the PGI is an important driver of the lack of concordance across PGIs, and that rigidly ranking individuals into quantiles based on current-day PGIs will inevitably lead to major misclassifications of true genetic risk.

## Results

### Rank concordance across PGIs

We present the results on the rank concordance of PGIs constructed with (i) different construction methods (i.e., C+T with *p* value threshold = 1, and LDpred with a prior fraction of causal SNPs = 1), but the same GWAS discovery sample; and (ii) the same construction method, but two different GWAS discovery samples (i.e., UKB and CARDIoGRAM/23andMe for CVD/EA respectively). The UKB GWAS discovery sample includes those of European ancestry and excludes the sibling holdout sample and their relatives. The subset of UKB siblings serves as our holdout sample here (Supplementary Information 1.1). Using bivariate LD Score regression^32^, we find that the genetic correlations between the GWAS summary statistics from different discovery samples are high. We estimate the genetic correlation *r*_g_ between the discovery samples to be 0.96 (SE = 0.03) for CVD and 0.88 (SE = 0.01) for EA. Hence, SNP effect sizes are generally concordant between the GWAS discovery samples. This suggests that differences in the environmental context of the discovery sample cannot be the main driver for discordance between PGIs, especially for CVD.

The LDpred PGI based on the meta-analysis of two samples (UKB and 23andMe for EA; UKB and CARDIoGRAM for CVD) results in the PGI with the highest explained variance for each trait (Fig. 1d and Fig. 2d). We refer to these PGIs as the “benchmark PGIs”. Fig. 1 and Fig. 2 visualise the rank concordance between the different PGIs for EA and CVD, respectively. Fig. 1a and Fig. 2a show the rank concordance in deciles of the PGI, with the size and shading of the bubble visualizing the extent of overlap. With full concordance of PGI deciles, all circles would fall on the diagonal line and be of the same size. However, we find especially low rank concordance between PGIs that use different GWAS discovery samples, and there is more discordance for PGIs constructed using C+T compared to those using LDpred. Fig. 1b and Fig. 2b show the Spearman correlation matrix for the differently constructed PGIs. We find rank correlations ranging between r = 0.40-0.85 for EA PGIs, and between r = 0.17-0.94 for CVD PGIs.

**Figure 1.**
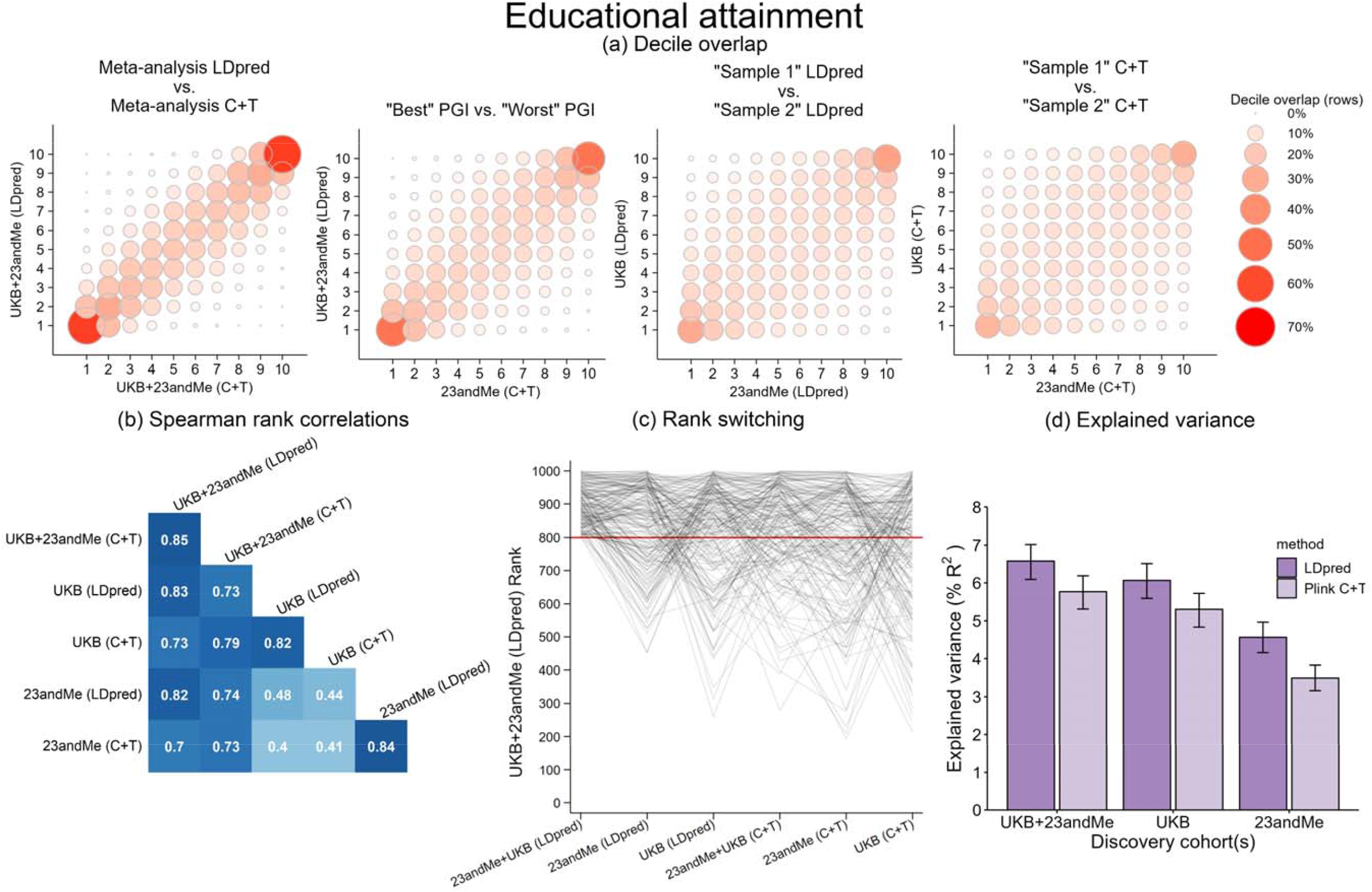
Concordance across six PGIs for educational attainment. **a**. Rank concordance in deciles of the PGI distribution. **b**. Spearman rank correlations across PGIs. **c**. rank switching across the PGIs for a random *N* = 1,000 individuals from the UKB holdout sample, with the red line denoting the top quintile. **d**. explained phenotypic variance of the PGIs with error bars showing 95% confidence intervals. PGI = polygenic index, C+T = clumping and thresholding.

**Figure 2.**
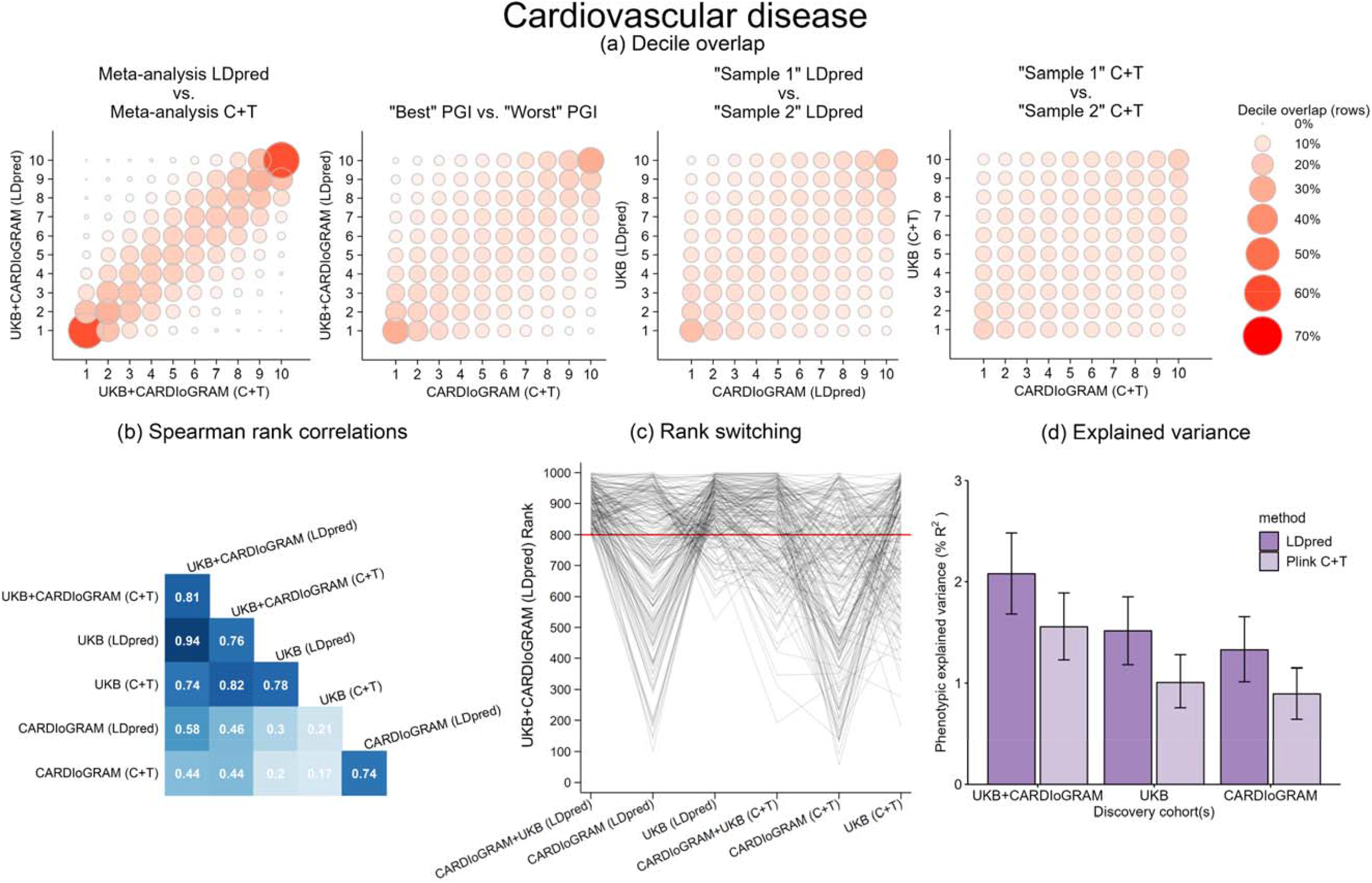
Concordance across six PGIs for cardiovascular disease. **a**. Rank concordance in deciles of the PGI distribution. **b**. Spearman rank correlations across PGIs. **c**. Rank switching across the PGIs for a random *N* = 1,000 individuals from the UKB holdout sample, with the red line denoting the top quintile. **d**. Explained phenotypic variance of the PGIs in terms of pseudo-*R*^2^ from a logit regression with error bars showing 95% confidence intervals. PGI = polygenic index, C+T = clumping and thresholding.

In Fig. 1c and Fig. 2c, the rank switching is visualised for a random subset of *N* = 1,000 individuals from our analysis sample. The vertical axis shows the exact PGI rank for these 1,000 individuals on the benchmark PGI, highlighting those in the top quintile (i.e., those above the red threshold line). The horizontal axis displays the different PGI construction methods. The lines show the extent to which individuals who are in the top quintile of the benchmark PGI switch ranks when using different construction methods. With full rank concordance, all “top”-ranked individuals would remain above the red line. However, we observe strikingly large rank switching: of all those in the top quintile of the “benchmark PGI” for CVD (CARDIoGRAM+UKB, LDpred), we find that between 21% (for our second-best performing PGI – CVD (UKB, LDpred)) and 63% (for our worst-performing PGI - CVD (CARDIoGRAM, C+T)) of individuals fall outside of the top quintile, i.e., move below the red line in panel (c). Only 10% of the individuals are in the top quintile for each of the six CVD PGIs. For EA, only 22% of individuals who are in the top quintile of the benchmark PGI are also in the top quintile of rest of the PGIs.

### Personalised interventions

We examine the extent to which rank switching between PGIs may influence individualised drug prescription for CVD. We illustrate the overlap in individuals to be prescribed statins (a type of cholesterol-lowering medication) according to recently proposed clinical guidelines to involve PGI data^22^. While statins reduce the risk of cardiovascular events in individuals with high cholesterol levels^33^, their benefits need to be assessed against their potential adverse effects, which includes a higher risk of developing diabetes^34^. Current guidelines from the American College of Cardiologists/American Heart Association (ACC/AHA) recommend statins for three groups of patients: those with high LDL cholesterol (≥190 mg/dL); 2) those with a combination of elevated LDL cholesterol (≥70 mg/dL) and diabetes; and 3) those with a combination of elevated LDL cholesterol (≥70 mg/dL) and a ≥7.5% (“high”) risk to develop atherosclerotic cardiovascular disease (ASCVD) within ten years^35^. These ten-year ASCVD risks can be calculated with prediction models from the ACC/AHA^36^. The ACC/AHA welcomes the inclusion of alternative risk factors to identify additional individuals who might benefit from statin therapy because they are at “borderline” (i.e., ≥5%) ten-year ASCVD risk^22^. Accordingly, an earlier study^22^ uses the top quintile of an LDpred PGI (based on CARDIoGRAM 2015 GWAS results^25^) as a risk factor to identify additional candidates for statin therapy for CVD-free individuals at borderline ASCVD risk. We follow their strategy and examine the variation in individuals to be recommended statins based on differentially constructed PGIs. For this analysis, we create a CVD-free holdout subsample in the UKB siblings sample (*N* = 4,061) consisting of individuals i) who report to not use statins and without a history of CVD; ii) who are not recommended statins according to current ACC/AHA guidelines; iii) who do have a have ≥5% ten-year ASCVD risk; and iv) who score in the top quintile of at least one of five CVD PGIs (we here drop the meta-analysed score from UKB + CARDIoGRAM (C+T) for visualisation purposes). The threshold determining the top PGI quintile is calculated in the full UKB holdout sample (i.e., including individuals who do have a history of CVD or use statins).

Fig. 3 shows a Venn diagram depicting the overlap in individuals’ ranking in the top quintile across the five CVD PGIs. Only 6% of the individuals are in the top quintile for all five CVD PGIs (inner cell), while 38% of the individuals are in the top quintile for only one PGI (outer layer). Discordance is especially high for PGIs created based on CARDIoGRAM GWAS summary statistics only (as in the study we follow here), with 35% (i.e., 10% + 12% + 13%) of the individuals scoring in the top quintile of the CARDIoGRAM C+T or LDpred PGI distributions but not in the other PGI distributions. Out of the *N* = 2,007 individuals eligible for statins according to the meta-analysis (UKB + CARDIoGRAM) LDpred PGI, only 38.6% would have received statins if this decision was based on the CARDIoGRAM (C+T) PGI instead. These results show that different sets of individuals may be selected for personalised intervention based on decisions made at the PGI construction stage.

**Figure 3.**
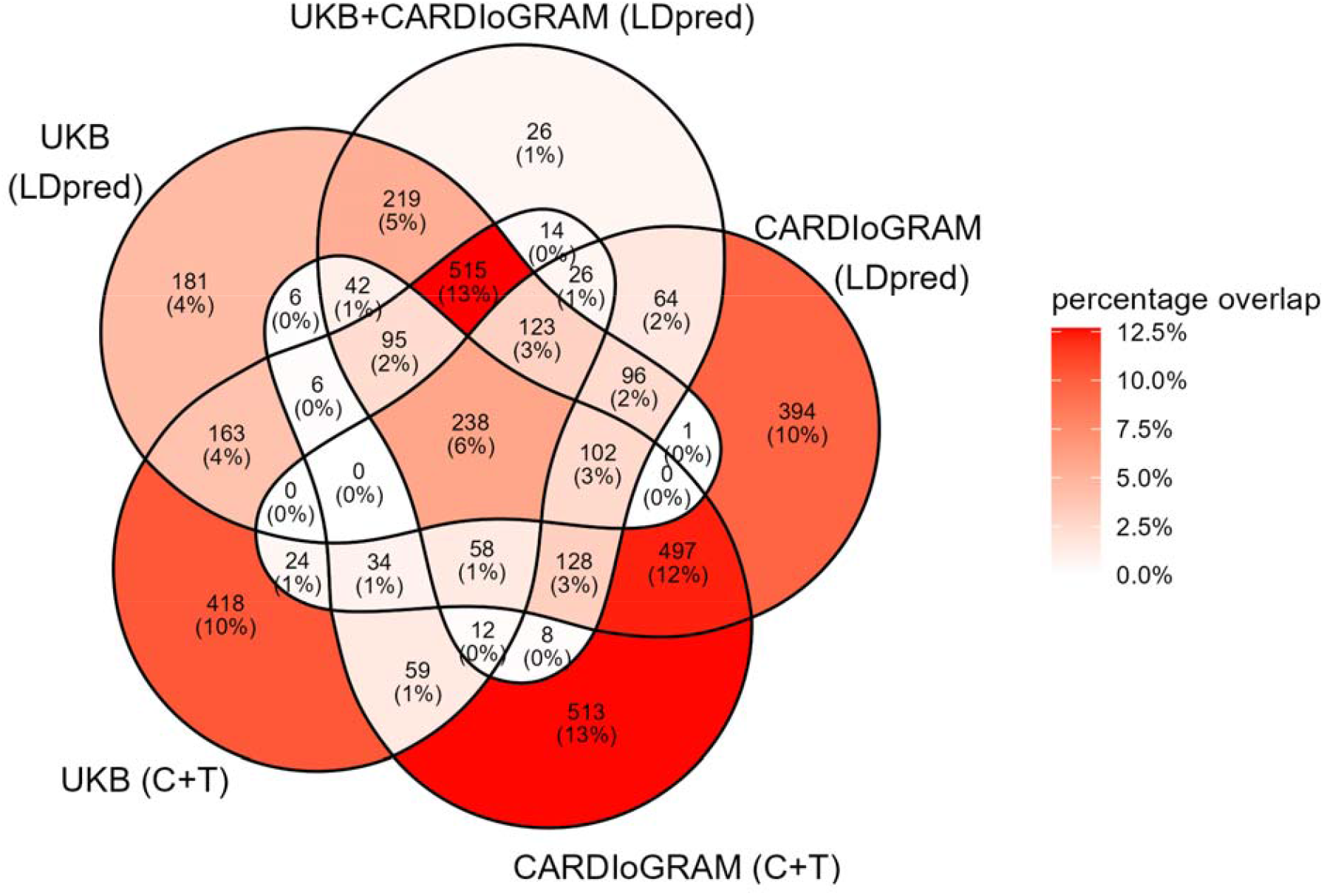
Venn diagram depicting the overlap in individuals ranked in the top quintiles of five CVD PGIs (*N* = 4,061). Individuals included in this figure are potential candidates for statin therapy : they have an intermediate ten-year ASCVD risk (≥5%); have no (self-reported) history of CVD; are not statin users; and are not yet candidates according to current ACC/AHA guidelines. C+T = clumping and thresholding.

### G×E interplay

We then explore whether the estimation of G×E interaction effects may vary by PGI construction method. G×E research explores how environments can moderate genetic susceptibilities, or vice versa, how genetic susceptibility can moderate environmental effects. For instance, assessing if the effectiveness of drug treatment varies by quantiles of genetic risk is a form of G×E research. If, however, the extent of genetic susceptibility in empirical studies is dependent on how the PGI is constructed, so may its estimated interaction with the environment. Here, we explore whether PGI rank discordance can affect G×E estimates. We follow a previous study design which found that association between the EA PGI and EA has decreased over time in the United States^31^, and explore how different methods of PGI construction influence the association between six different EA PGIs and EA across birth cohorts in the UKB.^31^

We assess whether modelling the PGI as a continuous or stratified measure of genetic predisposition alters the estimation of G×E. We run two regressions: in the first regression we use the EA PGI as a continuous variable, in the second regression we employ four binary indicators for the EA PGI quintiles. We do this separately for the six different PGIs. Fig. 4a shows the coefficients of the interaction term between the continuous PGI and year of birth, while Fig. 4b shows the coefficients of the interaction term between year of birth and PGI quintiles 1, 2, 4 and 5 (with quintile 3 serving as the baseline). In line with earlier evidence^31^, we find that the size of the association between the EA PGI and EA decreases for later-born cohorts as evidenced by the negative interaction terms in Fig. 4a and the negative slope of the interaction terms over the PGI quintiles in Fig. 4b. The most negative interaction terms in Fig. 4a are estimated using the two UKB-based PGIs, which is mirrored by the clearest negative gradient in Fig. 4b for the quintile-stratified PGIs. The interaction term that is closest to zero in Fig. 4a is estimated using the two 23andMe-based PGIs, which is mirrored by a less clear negative gradient in Fig. 4b. Although the estimates do not vary greatly across the different PGIs in that they are all negative and mostly statistically significant, a joint *F*-test rejects the hypothesis that the interaction coefficients are equal to each other. A pairwise *F*-test suggests that the divergence is driven by PGIs constructed using C+T (Supplementary Information 1.6). Overall, these results again illustrate that choices made at the PGI construction stage can affect the results in G×E analyses.

**Figure 4.**
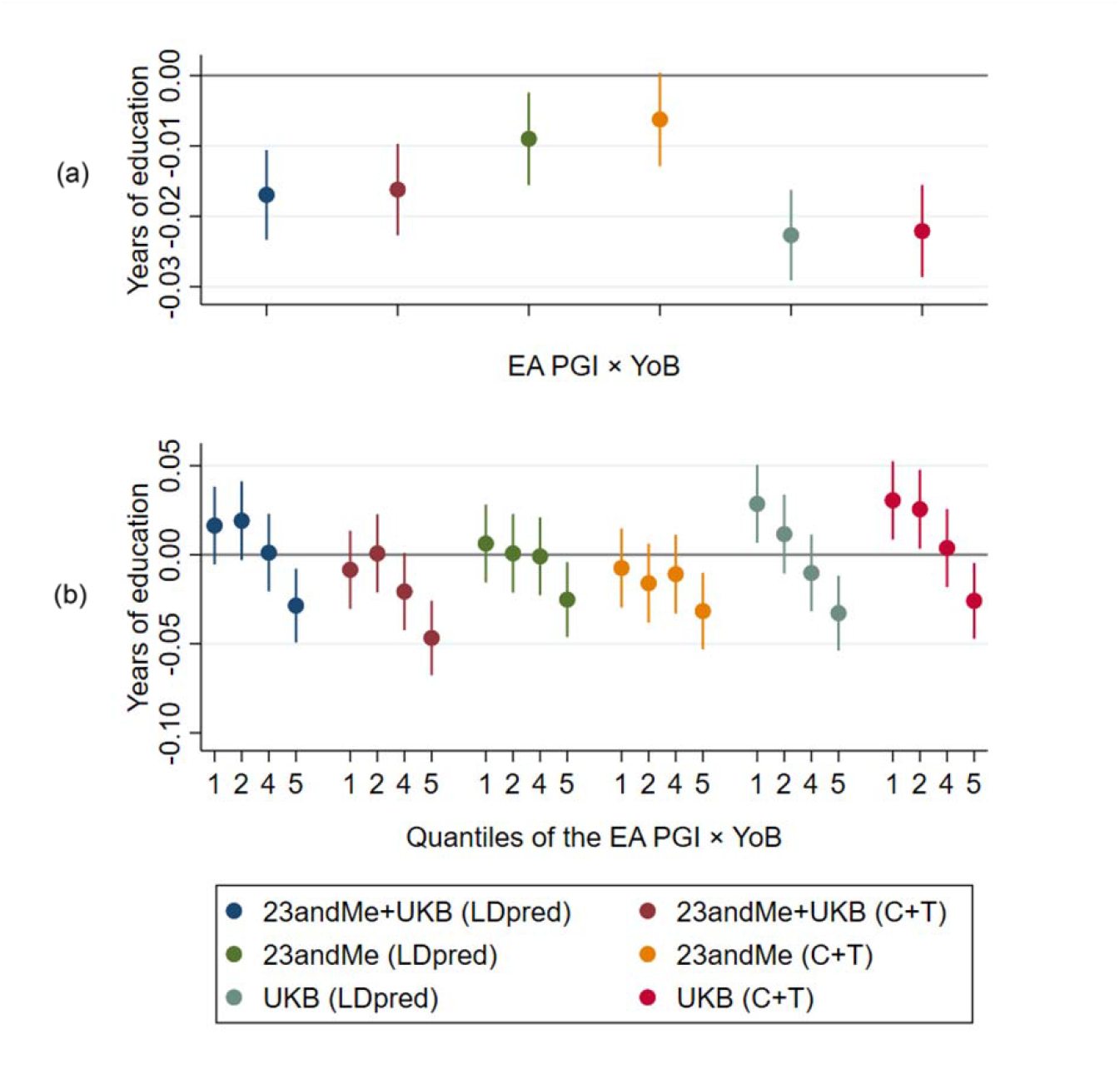
Results of OLS regressions explaining years of education by the EA polygenic index (PGI), year of birth (YoB), and the interaction between the EA PGI and YoB in the subsample of siblings of the UK Biobank (*N* = 38,049). **a**. The PGI analysed as a continuous variable. **b**. The PGI split into binary indicators for each quintile (with quintile 3 serving as the baseline). The figure visualises the estimated interaction terms with their 95% confidence intervals. C+T = clumping and thresholding.

### Simulations

The empirical analyses show that choices during the PGI construction phase may lead to the ranking of individuals in different quantiles of the resulting PGI distribution, but the underlying reason for such discordance across PGIs is not clear. Here, we use simulations to assess to what extent a discordance across different PGIs could be the result of measurement error in the PGI. Measurement error in the PGI stems from the fact that any underlying GWAS is conducted on a finite sample, with the coefficients that are used to construct the PGI exhibiting a degree of statistical noise that decreases with the size of the GWAS discovery sample^37^. As a result of measurement error in the coefficients, the predictive power of the PGI will fall short of the SNP-based heritability, which constitutes the upper bound of the predictive power of a PGI in terms of variance explained^38^. We define the “*explained* SNP-based heritability” as the ratio of the explained variance of a PGI and the SNP-based heritability of the trait of interest. In our simulations, we model measurement error to be classical, and we use EA as our benchmark trait, which has a SNP-heritability of around 25%, a PGI that explains 12% of its variation, and with that, an *explained* SNP-heritability of around 50%^39,40^.

Fig. 5 shows that “rank precision” of a PGI (i.e., the fraction of individuals correctly classified into the top quintile of the “true” PGI) strongly depends on the *explained* SNP-based heritability. Naturally, an explained SNP-based heritability of 100% is necessary to accurately place individuals in the top quintile of the PGI distribution. For 80% accuracy, an explained SNP-based heritability of 88% is needed. With a current explained SNP-based heritability of 50% for EA, we can expect a 57% correct placement in the top quintile of the PGI distribution.

**Figure 5.**
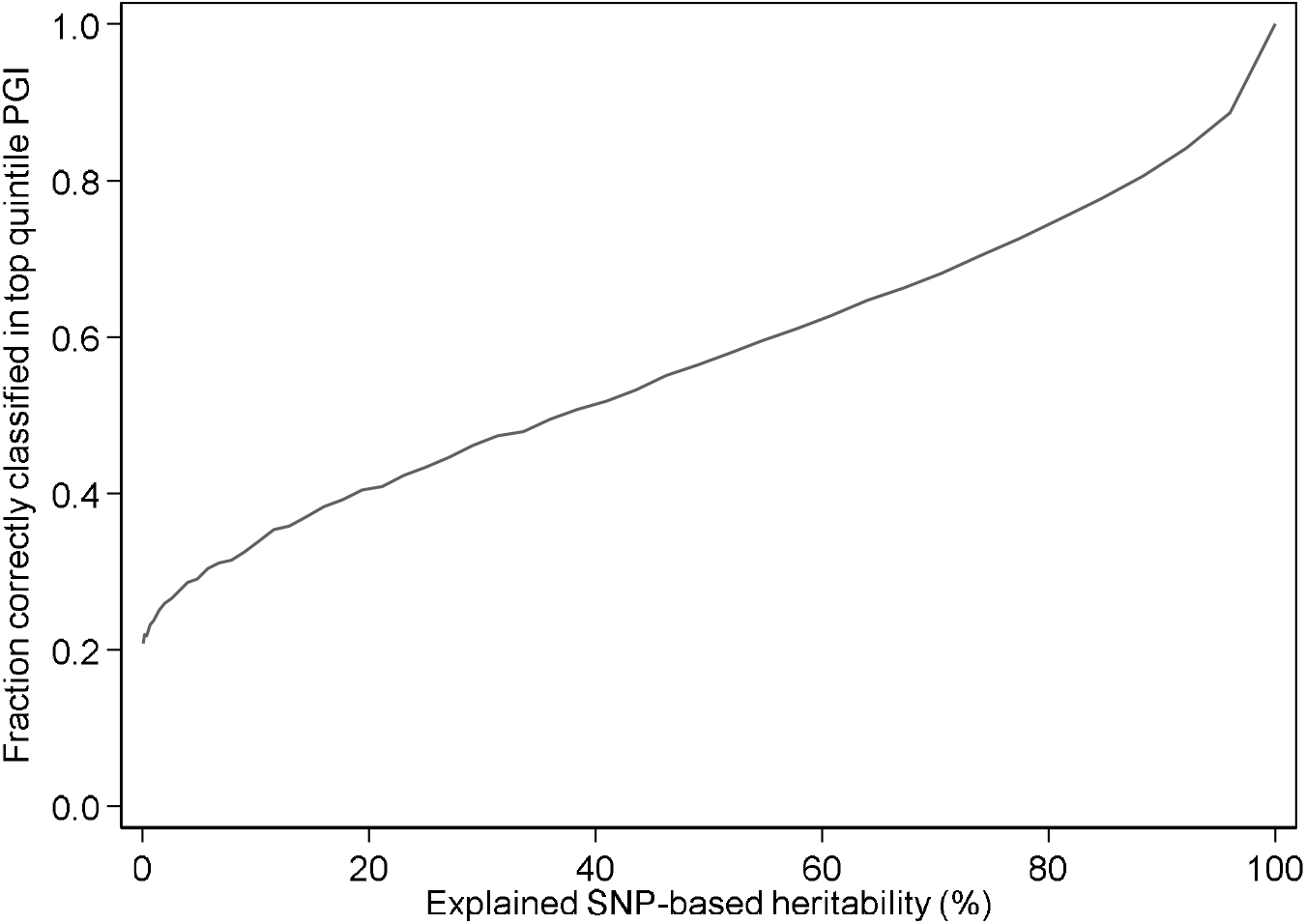
The relationship between the predictive power of the PGI and the correct classification of individuals in the top quintile of the PGI distribution. This figure visualises the relationship between the explained SNP-based heritability (%) and the fraction of correctly classified individuals in the top quintile of the PGI distribution.

Fig. 6 shows decile overlap between the simulated “true” PGI and PGIs with varying degrees of explained SNP-based heritability, quantifying to which extent individuals are placed into the correct decile of the true PGI given a noisy PGI. While concordance increases with increasing explained SNP-based heritability, even for a PGI with an explained SNP-based heritability of 80% the fraction of off-diagonal elements is only 72.7 percent.

**Figure 6.**
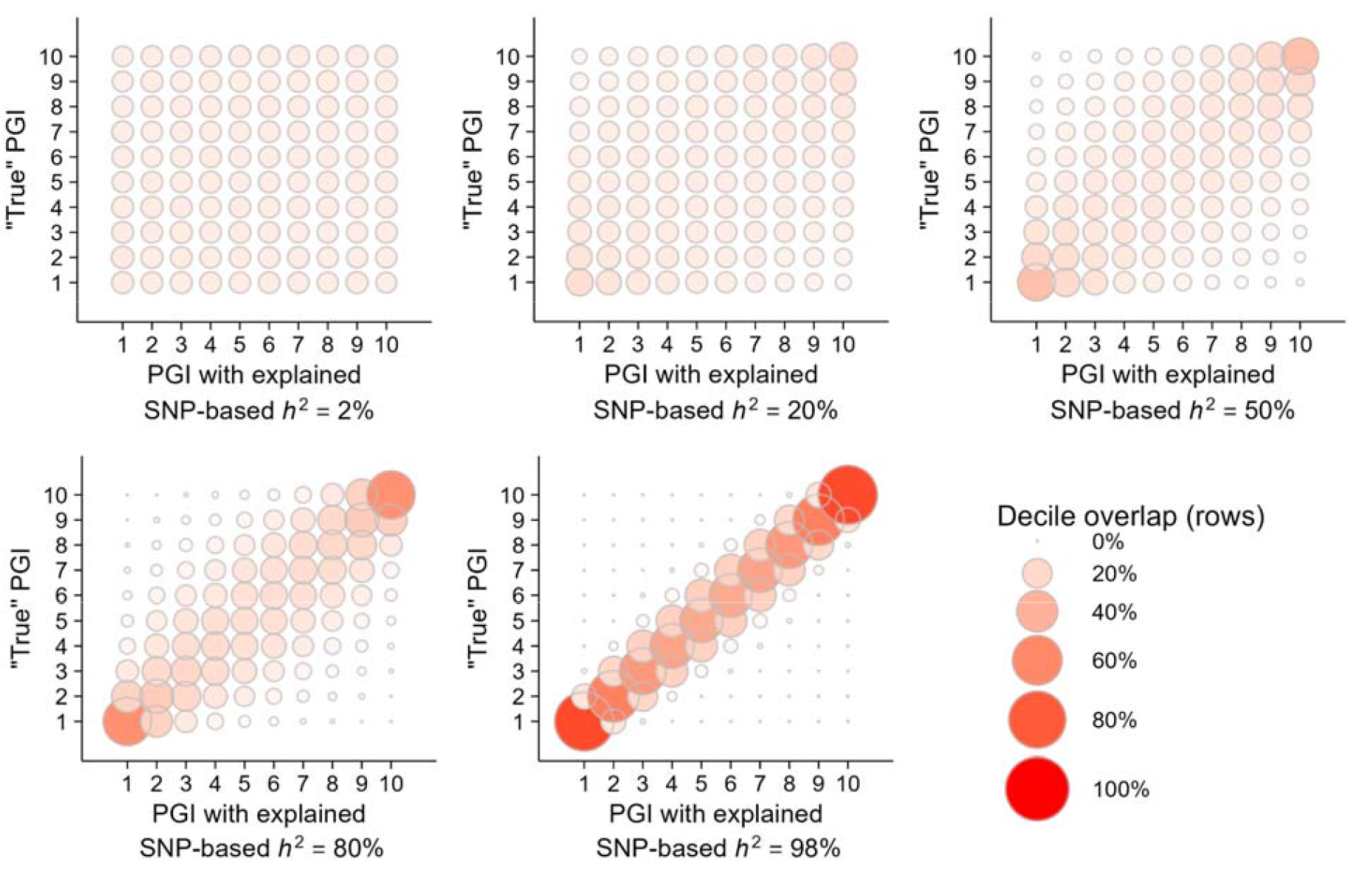
Results of the simulations analysing the relationship between the predictive power of a PGI and the ranking of individuals in the PGI distribution. This figure shows the rank concordance in terms of deciles between the “true” PGI and PGIs with varying degrees of explained SNP-based heritability (*h*^*2*^).

In Supplementary Information 1.7, we show that these simulation results are independent of the absolute level of the SNP-based heritability of a trait. In other words, correct classification into quintiles of genetic risk depends on the *explained* SNP-based heritability, regardless of the absolute level of heritability. Correct classification into the quintiles of the *trait* distribution, however, does depend on the absolute level of the SNP-based heritability. For example, the 25% SNP-based heritability for EA implies that, even with a perfect PGI, the prediction accuracy of the correct quintile in the *trait* distribution is only around 40% (Supplementary Information 1.7). Thus, the increasing predictive power of PGIs implies better rank concordance across PGIs due to increases in the explained SNP-based heritability. Nonetheless, prediction accuracy of the PGI on the trait level is constrained by the SNP-based heritability of the trait. Hence, there are two layers of uncertainty when using PGIs for trait prediction: first, any estimated PGI is a noisy proxy for the “true” PGI, and second, any risk prediction of even the “true” PGI is limited by the SNP-based heritability.

## Discussion

Despite high genetic correlations between GWAS discovery samples, the ranking of individuals across differently constructed PGIs can vary substantially. This rank discordance between PGIs can have implications for personalised interventions and gene-environment interaction research. We focus on two traits that have recently garnered attention as candidates for individualised intervention: cardiovascular disease (CVD) for individualised drug prescription, and educational attainment (EA) for individualised learning trajectories. PGIs for both traits are also currently being put to use for pre-implantation genetic testing for embryos^9^.

We show that using differentially constructed CVD PGIs for individualised statin prescription identifies different groups of individuals eligible for statins, with only 6% of individuals in our sample ranking consistently in the top quintile for each of the five PGIs (Fig. 3). Importantly, misclassifications may lead to adverse treatment effects^34^. With regards to educational attainment, our simulations show that we classify just over 50% of individuals correctly in the top quintile of the “true” PGI with current-day PGIs. Hence, using PGIs for “precision education”^11^ is likely to lead to educational customisations that are channelled to the wrong individuals in a substantial number of cases. Classifying individuals into quantiles of a PGI distribution can also have repercussions for empirical research, as we find that the PGI construction method can affect the estimates of the importance of the nature-nurture interplay in shaping life outcomes.

Our study joins earlier studies in their call for making the use and reporting of PGIs and their construction more transparent and standardised^17–19,41^ and contributes to the set of recent studies highlighting the divergent predictive power of PGIs^20,42–44^. Pain et al. compare a very extensive set of traits and test the predictive power of a wide variety of PGI construction methods^43^. Ware et al. compare a more limited set of PGI construction methods and analyse the intra-individual correlation of PGIs^42^. Finally, two studies that were independently developed around the same time^20,44^ are similar in spirit as the present study in comparing the rank concordance of individuals in the PGI distribution depending on the GWAS discovery sample. Our study complement these studies by i) explicitly focusing on rank discordance and its source, ii) comparing across PGI construction methods (e.g., C+T and LDpred), and iii) analysing the implications for empirical applications such as personalised medicine or G×E analysis.

Our findings are of crucial importance now that PGIs are becoming increasingly accessible to physicians, consumers, and applied researchers^19^. We complement recent work that showed that an individual’s PGI can span several deciles when the uncertainty of GWAS estimates are taken into account during PGI construction^45,46^. While the source of uncertainty emphasised in these papers does not derive from the construction method or GWAS discovery sample per se, we draw a similar conclusion: ranking individuals on basis of their position in a PGI distribution is prone to large uncertainty. Therefore, transparent reporting^17^ and robustness checks against different PGIs should become routine in analyses that use PGI ranks. We conclude that while PGIs can be a useful tool for identifying individuals at risk, rigidly relying on a PGI rank from a single (noisy) PGI may lead to misinformed decision and policymaking.

## Methods

### Sample and data

Participants of this study were sourced from UK Biobank, a prospective cohort study in the UK that collects physical, health and cognitive measures, and biological samples (including genotype data) in about 500,000 individuals^24^. UK Biobank has received ethical approval from the National Health Service North West Centre for Research Ethics Committee (11/NW/0382) and has obtained informed consent from its research participants. In our analyses, we include only European ancestry respondents (81% of the UKB). The UK Biobank’s sibling subsample serves as the holdout sample. Siblings and their relatives are identified using the UKB’s kinship matrix based on genetic relatedness and containing relatives of third degree and closer. The sibling subsample consists of *N* = 39,296 individuals (16,556 males and 22,740 females). The age of these individuals ranges from 40 to 71 years with the average age of 57. years at recruitment. More information about the analysis sample and the construction of variables can be found in Supplementary Information 1.1 and 1.2.

### Statistical analyses

The PGIs used in this study are based on four sets of GWAS summary statistics: GWAS summary statistics for EA (*N* = 389,419) and CVD (*N* = 392,789) resulting from our GWAS conducted in the UKB sample (excluding the siblings subsample and their relatives, Supplementary Information 1.3), GWAS summary statistics for EA from 23andMe (*N* = 365,536), and GWAS summary statistics for CVD from the CARDIoGRAM^25^ consortium (*N* = 184,305). Mixed linear model GWAS were conducted with fastgwa^47^, using the sparse genotype matrix provided by the UKB to account for relatedness between participants. The CVD phenotype is based on hospital and death records (ICD9 410-414 or ICD10 I20-I25). The PGIs were constructed using LDpred^29^ (prior fraction of causal SNPs = 1) and Plink clumping and thresholding^28^ (*p* value threshold is = 1, more details in Supplementary Information 1.4). The personalised intervention analysis (Supplementary Information 1.5) and G×E analyses (Supplementary Information 1.6) as well as the simulations (Supplementary Information 1.7) were performed in STATA.

## Data availability statement

Individual-level genotype and phenotype data are available by application via the UKB Biobank website (https://www.ukbiobank.ac.uk/). The genome-wide summary statistics from 23andMe can be obtained by completing the 23andMe publication dataset access request form at https://research.23andme.com/dataset-access/. The genome wide summary statistics from CARDIoGRAM are available at http://www.cardiogramplusc4d.org/. The authors declare that the results supporting the findings of this study are available within the paper and its supplementary information files.

## Code availability statement

Analysis code will be made available on Github.

## Acknowledgments

UK Biobank has obtained ethical approval from the National Research Ethics Committee (11/NW/0382). This research has been conducted using the UK Biobank Resource under application number 41382. The authors gratefully acknowledge funding from NORFACE through the Dynamic of Inequality across the Life Course (DIAL) programme (GEIGHEI 462-16-100). Research reported in this publication was also supported by the National Institute on Aging of the National Institutes of Health under Award R56AG058726. S.F.W.M. gratefully acknowledges funding from the European Union’s Horizon 2020 research and innovation program under the Marie Skłodowska-Curie grant agreement (GENIO 101019584). C.A.R. and S.v.H. gratefully acknowledge funding from the European Research Council (GEPSI 946647; DONNI 851725). We are grateful for Aysu Okbay and employees and research participants of the 23andMe, Inc. cohort for sharing GWAS summary statistics for educational attainment, and we thank Pietro Biroli, Titus Galama, and Eric Slob for insightful comments. This work made use of the Dutch national e-infrastructure with the support of the SURF Cooperative using grant EINF-1107.

## Author contributions

S.F.W.M. and D.M. designed and oversaw the study. S.F.W.M. and D.M. conducted the GWAS in UKB and the meta-analyses with other GWAS summary statistics, constructed the PGIs, and prepared the illustrative applications. R.D.P. performed the G×E analyses. H.v.K. conducted the simulations. C.A.R. and S.v.H. assisted with the analyses. All authors contributed to preparing and critically reviewing the manuscript and the supplementary file.

## References

1. Visscher, P. M. et al. 10 years of GWAS discovery: biology, function, and translation. Am. J. Hum. Genet. 101, 5–22 (2017).

2. Chabris, C. F., Lee, J. J., Cesarini, D., Benjamin, D. J. & Laibson, D. I. The fourth law of behavior genetics. Curr. Dir. Psychol. Sci. 24, 304–312 (2015).

3. Dudbridge, F. Power and predictive accuracy of polygenic risk scores. PLoS Genet. 9, 1003348 (2013).

4. The International Schizophrenia Consortium. Common polygenic variation contributes to risk of schizophrenia and bipolar disorder. Nat. Lett. 460, 748–752 (2009).

5. Khera, A. V. et al. Genome-wide polygenic scores for common diseases identify individuals with risk equivalent to monogenic mutations. Nat. Genet. 50, 1219–1224 (2018).

6. Mega, J. L. et al. Genetic risk, coronary heart disease events, and the clinical benefit of statin therapy: an analysis of primary and secondary prevention trials. Lancet 385, 2264–2271 (2015).

7. Torkamani, A., Wineinger, N. E. & Topol, E. J. The personal and clinical utility of polygenic risk scores. Nat. Rev. Genet. 19, 581–590 (2018).

8. Kumar, A. et al. Whole-genome risk prediction of common diseases in human preimplantation embryos. Nat. Med. 28, 513–516 (2022).

9. Turley, P. et al. Problems with using polygenic scores to select embryos. N. Engl. J. Med. 385, 79– 85 (2021).

10. Johnston, J. & Matthews, L. J. Polygenic embryo testing: understated ethics, unclear utility. Nat. Med. 28, 446–448 (2022).

11. Von Stumm, S. & Plomin, R. Using DNA to predict intelligence. Intelligence 86, 101530 (2021).

12. Shero, J. et al. The practical utility of genetic screening in school settings. npj Sci. Learn. 6, 1–10 (2021).

13. Biroli, P. et al. The economics and econometrics of gene-environment interplay. ArXiv (2022) doi:10.48550/arXiv.2203.00729.

14. Pereira, R. D., van Kippersluis, H. & Rietveld, C. A. The interplay between maternal smoking and genes in offspring birth weight. MedRxiv (2020) doi:10.1101/2020.10.30.20222844.

15. Barcellos, S. H., Carvalho, L. S. & Turley, P. Education can reduce health differences related to genetic risk of obesity. Proc. Natl. Acad. Sci. U. S. A. 115, E9765–E9772 (2018).

16. Slob, E. A. W. & Rietveld, C. A. Genetic predispositions moderate the effectiveness of tobacco excise taxes. PLoS One 16, e0259210 (2021).

17. Wand, H. et al. Improving reporting standards for polygenic scores in risk prediction studies. Nature 591, 211–219 (2021).

18. Lambert, S. A. et al. The polygenic score catalog as an open database for reproducibility and systematic evaluation. Nat. Genet. 53, 420–425 (2021).

19. Becker, J. et al. Resource profile and user guide of the polygenic index repository. Nat. Hum. Behav. 5, 1744–1758 (2021).

20. Schultz, L. M. et al. Stability of polygenic scores across discovery genome-wide association studies. BioRxiv (2021) doi:https://doi.org/10.1101/2021.06.18.449060.

21. Mills, M. C., Barban, N. & Tropf, F. C. An introduction to statistical genetic data analysis. Cambridge MIT Press. (2020).

22. Aragam, K. G. et al. Limitations of contemporary guidelines for managing patients at high genetic risk of coronary artery disease. J. Am. Coll. Cardiol. 75, 2769–2780 (2020).

23. Inouye, M. et al. Genomic risk prediction of coronary artery disease in 480,000 adults: implications for primary prevention. J. Am. Coll. Cardiol. 72, 1883–1893 (2018).

24. Bycroft, C. et al. The UK Biobank resource with deep phenotyping and genomic data. Nature 562, 203–209 (2018).

25. Nikpay, M. et al. A comprehensive 1,000 genomes-based genome-wide association meta-analysis of coronary artery disease. Nat. Genet. 47, 1121–1130 (2015).

26. Lee, J. J. et al. Gene discovery and polygenic prediction from a genome-wide association study of educational attainment in 1.1 million individuals. Nat. Genet. 50, 1112–1121 (2018).

27. Purcell, S. et al. PLINK: a tool set for whole-genome association and population-based linkage analyses. Am. J. Hum. Genet. 81, 559–575 (2007).

28. Chang, C. C. et al. Second-generation PLINK: rising to the challenge of larger and richer datasets. Gigascience 4, 1–16 (2015).

29. Vilhjálmsson, B. J. et al. Modeling linkage disequilibrium increases accuracy of polygenic risk scores. Am. J. Hum. Genet. 97, 576–592 (2015).

30. Ni, G. et al. A comparison of ten polygenic score methods for psychiatric disorders applied across multiple cohorts. MedRxiv (2021) doi:10.1101/2020.09.10.20192310.

31. Conley, D., Laidley, T. M., Boardman, J. D. & Domingue, B. W. Changing polygenic penetrance on phenotypes in the 20th century among adults in the US population. Sci. Rep. 6, 6–10 (2016).

32. Yengo, L., Yang, J. & Visscher, P. M. Expectation of the intercept from bivariate LD score regression in the presence of population stratification. BioRxiv (2018) doi:10.1101/310565.

33. Adhyaru, B. B. & Jacobson, T. A. Safety and efficacy of statin therapy. Nat. Rev. Cardiol. 15, 757– 769 (2018).

34. Torkamani, A., Wineinger, N. E. & Topol, E. J. The personal and clinical utility of polygenic risk scores. Nat. Rev. Genet. 19, 581–590 (2018).

35. Grundy, S. M. et al. 2018 AHA/ACC/AACVPR/AAPA/ABC/ACPM/ADA/AGS/APhA/ASPC/NLA/PCNA guideline on the management of blood cholesterol: executive summary. J. Am. Coll. Cardiol. 73, 3168–3209 (2019).

36. Goff, D. C. et al. 2013 ACC/AHA guideline on the assessment of cardiovascular risk: a report of the american college of cardiology/american heart association task force on practice guidelines. J. Am. Coll. Cardiol. 63, 2935–2959 (2014).

37. Van Kippersluis, H. et al. Using obviously-related instrumental variables to increase the predictive power of polygenic scores. BioRxiv (2020) doi:10.1101/2021.04.09.439157.

38. Witte, J. S., Visscher, P. M. & Wray, N. R. The contribution of genetic variants to disease depends on the ruler. Nat. Rev. Genet. 5, 765–776 (2014).

39. Okbay, A. et al. Genome-wide association study identifies 74 loci associated with educational attainment. Nature 533, 539–542 (2016).

40. Tropf, F. C. et al. Hidden heritability due to heterogeneity across seven populations. Nat. Hum. Behav. 1, 757–765 (2017).

41. Choi, S. W., Mak, T.S.-H. & O’Reilly, P. F. Tutorial: a guide to performing polygenic risk score analyses. Nat. Protoc. 15, 2759–2772 (2020).

42. Ware, E. B. et al. Heterogeneity in polygenic scores for common human traits. BioRxiv (2017) doi:10.1101/106062.

43. Pain, O. et al. Evaluation of polygenic prediction methodology within a reference-standardized framework. PLoS Genet. 17, 1–22 (2021).

44. Clifton, L., Collister, J. A., Liu, X., Littlejohn, T. J. & Hunter, D. J. Assessing agreement between different polygenic risk scores in the UK Biobank. MedRxiv (2022) doi:10.1101/2022.02.09.22270719.

45. Sun, J. et al. Translating polygenic risk scores for clinical use by estimating the confidence bounds of risk prediction. Nat. Commun. 12, 1–9 (2021).

46. Ding, Y. et al. Large uncertainty in individual polygenic risk score estimation impacts PRS-based risk stratification. Nat. Genet. 54, 30–39 (2022).

47. Jiang, L. et al. A resource-efficient tool for mixed model association analysis of large-scale data. Nat. Genet. 51, 1749–1755 (2019).

