## Supplementary Information for "Rank concordance of polygenic indices: Implications for personalised intervention and gene-environment interplay"

### 1. Data & Methods

#### 1.1. UK Biobank

The UK Biobank (UKB) provides genetic and socio-economic information for 500,000 volunteering and consented individuals^1^. Participation in the UK Biobank is voluntary, hence, it is not a representative sample of the UK population. The UKB cohort consists of an adult population, 40 to 70 years old between 2006-2010^2^. We include only European ancestry respondents (94%).

We use the UK Biobank’s sibling sub sample as the main holdout. We identify siblings and their relatives using the UKB’s kinship matrix based on genetic relatedness and containing relatives of third degree and closer. The matrix is computed using the KING software^3^. The UK Biobank does not have information about self-reported relatedness^2^. The degree of relatedness between the pairs of individuals is based on the combination of the kinship coefficient and genetic similarity in terms of the identity by state (IBS_0_) coefficient. IBS_0_ measures the fraction of markers for which the related individuals do not share alleles. We follow the KING manual for the thresholds on classifying the family relationships. For our analyses, we further separate those who are related to the siblings up to the 3rd degree (kinship coefficient ≥ 0.025), i.e., siblings, parents of siblings, cousins of siblings (Supplementary Table **1**). This ensures that our holdout sample for PGI construction and prediction (i.e., the sibling subsample) is unrelated to the GWAS discovery sample which is used to calibrate the SNP weights and construct the polygenic indices.

| Relationship to siblings | Unrelated to siblings | Full siblings | 2^nd^-3^rd^ relative of siblings | Parent or child of siblings | Total |
| --- | --- | --- | --- | --- | --- |
| *N* (individuals) | 91,055 | 41,498 | 10,207 | 4,740 | 147,500 |

**Supplementary Table 1.** Relatedness to the individuals in the siblings’ subsample of UK Biobank.

#### 1.2. Traits

We focus on two main phenotypes, outcomes, or traits to capture the differences by polygenicity of the traits and their potential use in gene-environment interaction studies or personalised intervention: educational attainment (EA) and cardiovascular disease (CVD).

We convert individuals’ highest earned qualifications to equivalent years of education using the International Standard Classification of Education (ISCED) following the literature^4–6^ to keep the outcome variables harmonized with those used in external GWAS such as e.g. 23andMe. Specifically, years of education ranges from 7 to 20, where College or University degree is equivalent to 20 years, National Vocational Qualification (NVQ), Higher National Diploma (HND), or Higher National Certificate (HNC) to 19 years, other professional qualifications to 15 years, having an A or AS levels similar to 13 years, O levels, (General) Certificate of Secondary Education ((G)CSE) to 10 years, and if none of the above to the lowest level of 7 years.

For cardiovascular disease (CVD), we focus on hospital admission and death records, ICD9 and ICD10 coding: ICD9 410-414 or ICD10 I20-I25. Thus, we define a “soft CVD” phenotype that includes angina in addition to myocardial infarction. We identify *N_cases_* = 29,970 and *N_controls_* = 362,819, and treat CVD as a linear phenotype in the fastGWA package, where the binary CVD phenotype is residualised on control covariates before we perform GWAS (Section 1.3).

#### 1.3. Genome-wide association analysis

For genome-wide association analysis (GWAS) we use the fastGWA tool for Genome-wide Complex Trait Analysis (GCTA)^7^, which applies mixed linear modelling (MLM) to the genetic data. fastGWA requires the following steps. First, we retain related individuals, generating a sparse genetic relatedness matrix (GRM) based on the family relatedness provided by the UKB. Next, we perform an MLM-based GWAS using the SNP data, the sparse GRM, the trait of phenotype of interest and the minor allele frequency (MAF) filter of 0.001. The phenotype is residualised with respect to birth year dummies, gender, interaction of birth year and gender, batch, and the first 40 principal components (PCs) of the genetic relatedness matrix. For quality control reasons, some individuals were not included in the GWAS. More specifically, we exclude individuals have missing gender or whose self-reported gender does not match the genetic sex, are of other than European ancestry, have poor genotyping quality, putative sex chromosome aneuploidy, whose second chromosome karyotypes are different from XX or XY, with outliers in heterozygosity, or have missing information on any of the former criteria. We also drop those who withdrew consent.

We meta-analyse our UKB GWAS with: GWAS from the CARDIoGRAM^8^ for CVD; GWAS from 23andMe^5^ for EA. We apply quality control to the resulting GWAS summary statistics using EasyQC^9^. Meta-analysis is conducted using the software package METAL^10^. The GWAS discovery sample sizes are reported in Supplementary Table 2.

#### 1.4. Polygenic index (PGI) construction

A PGI is a way of summarizing the effects of multiple genetic variants into a single score, and can be seen as the best linear genetic predictor of a phenotype^11,12^. For this reason, it was recently termed the “additive genetic factor”^13^. It is calculated as

${PGI}_{i}=\sum_{j=1}^{M} w_{j}a_{ij}$	(1)

where $a_{ij}$denotes the number of ‘effect’ alleles (0, 1, or 2) for each individual $i$ at Single-Nucleotide Polymorphism (SNP) $j$, $w_{j}$ denotes the weight given to the SNP from an independent GWAS, and *M* is the number of SNPs included^14^. Weights $w_{j}$ are equal to the coefficient of the estimated relationship between SNP *j* and the outcome variables in a GWAS. This continuous genetic score has many potential applications – for example as control variable in observational studies^15,16^, to boost power of randomized controlled trials (RCTs)^6^, as instrumental variables to establish causal effects of one phenotype on an outcome of interest^17^, to study interactions between genes and environmental exposures, and personalized medicine and pharmacogenetics.^[1]^

As is clear from equation (1), there are two basic choices to be made when constructing a PGI^12^:

1. Which SNPs to include?
2. What SNP weights to use?

There are two main approaches to select SNPs and their weights in constructing PGIs. The first one is known as ‘clumping and thresholding’ (C+T)^18^ as implemented in the software package Plink^19^. This method selects one SNP from a block of SNPs that is in linkage disequilibrium (LD, i.e., correlation across SNPs resulting from the fact that SNPs physically close within the same chromosome are often transmitted together). The selected SNP within each LD-block is either random (‘pruning’) or the SNP with the lowest *p* value is selected (‘clumping’). Maintaining the independent SNPs, the coefficient from the GWAS is then simply used as weight $w_{j}$ in equation 1. The second method retains all available HapMap3 SNPs that passed quality controland uses the correlation structure of these SNPs in a Bayesian approach to adjust the GWAS coefficients into weights $w_{j}$. The most commonly used method of this kind is LDpred^20,21^.

The first main contrast we will investigate is between polygenic indices constructed on basis of the same GWAS summary statistics using (i) Plink’s C+T algorithm and (ii) LDpred. The baseline comparison will be between a C+T score with a *p* value threshold of 1 (i.e., retaining a SNP for each LD block) and the LDpred with a prior of 1 (i.e., the assumed causal fraction of SNPs). LDpred^20^, version 1.06, and Python, version 3.6.6.) is a software package based on Python that adjusts the GWAS weights for linkage disequilibrium using a Bayesian approach. We follow the steps as outlined in a recent textbook on statistical genetics^12^, including the coordination of the base and target files, computing the LD adjusted weights, and then applying them for polygenic index construction using Plink^22^. We re-weight the SNP effects on the basis of LD and the supposed fraction of causal SNPs, which we set to 1, as is standard practice for highly polygenic traits^13^. Our hold-out sample for constructing polygenic indices consists of *N* = 39,326 siblings. At the coordination step, we filter for HapMap3 SNPs. The final number of SNPs remaining after filtering and quality control used for each polygenic index (PGI) is reported in Supplementary Table 2 below.

| Trait | Discovery Sample | Construction  Method | Discovery Sample Size | Number of SNPs | Incremental *R*^2^** |
| --- | --- | --- | --- | --- | --- |
| Educational Attainment | UKB* | LDpred | *N* = 389,419 | 1,065,147 | 6.1% |
|  |  | Plink C+T |  | 1,127,29 | 5.3% |
|  | 23andMe^5^ | LDpred | *N =* 365,536 | 1,057,144 | 4.6% |
|  |  | Plink C+T |  | 109,265 | 3.5% |
|  | Meta-analysis | LDpred | *N =* 754,955 | 1,065,061 | 6.6% |
|  |  | Plink C+T |  | 112,576 | 5.8% |
| Cardiovascular disease | UKB* | LDpred | *N*_Cases_ = 29,970  *N*_Controls_ = 362,819 | 1,065,145 | 1.5% |
|  |  | Plink C+T |  | 112,793 | 1.0% |
|  | CARDIoGRAM^8^ | LDpred | *N*_Cases_ = 60,801  *N*_Controls_ = 123,504 | 1,062,642 | 1.3% |
|  |  | Plink C+T |  | 110,439 | 0.9% |
|  | Meta-analysis | LDpred | *N*_Cases_ = 90,771  *N*_Controls_ = 486,323 | 1,065,062 | 2.1% |
|  |  | Plink C+T |  | 112,810 | 1.6% |

**Supplementary Table 2**: Overview of PGIs used in the study. Note: * UKB discovery sample always excludes siblings and their relatives up to a 3^rd^ degree (cousins). **Additional explained phenotypic variance after controlling for gender, genotyping batch, dummies for year and month of birth, interactions between gender and year of birth, and the first 40 principal components of the genetic relatedness matrix (holdout sample: *N* = 39,296 UKB siblings). For CVD, this is the pseudo-*R^2^* from a logit regression. CVD = cardiovascular disease. C+T = clumping and thresholding. Prior for LDpred = 1, *p* value threshold for C+T = 1

#### 1.5. Personalised intervention

In this analysis, we identify individuals in the holdout sample who are at borderline risk of developing a atherosclerotic cardiovascular event (ASCVD) in the next 10 years (10-year ASCVD risk ≥5%), and assess the utility of PGIs for CVD to expand recommendation guidelines^23^. We calculate ten-year ASCVD risk with the AHA Pooled Cohort Equations (PCE) for individuals of European ancestry^24^. This calculation is based on sex, age, total and HDL cholesterol levels, systolic blood pressure, smoking status, and diabetes. Individuals are recommended statins if they meet one of the following criteria: 1) LDL cholesterol ≥190 mg/dL; 2) LDL cholesterol ≥70 mg/dL and diabetes; 3) LDL cholesterol ≥70 mg/dL and 10-year risk of developing atherosclerotic cardiovascular disease (ASCVD) ≥7.5%. In accordance with earlier work^25^, we define diabetes using the self-reported diabetes status; self-reported insulin use; or a casual blood glucose level of ≥11 mmol/L. In our UKB holdout sample of siblings, we find that 9.7% (*N* = 2,484) of individuals (who are not currently using statins) should be prescribed statins according to ACC/AHA guidelines, and that the average 10-year ASCVD risk in CVD-free, non-statin using individuals is 5.5% (range: 4.5% - 47.2%). Similar to Aragam et al. (2020)^23^, we then use the different CVD PGIs to expand the recommendation of statin therapy to those in in the top quintile of a CVD PGI and at borderline to intermediate risk of ASCVD (i.e., 10-year ASCVD risk ≥5%). The CVD PGI quintiles are calculated based on the full holdout sample to capture the full spectrum of genetic risk, i.e., including those with CVD and statin use.

For this analysis, we remove individuals in the discovery sample or related to individuals in the discovery sample; those already prescribed statins according to self-report; and those already suffering from CVD, as defined by self-reported angina, myocardial infarction, or cardiovascular operations, and from hospital admission and death records (ICD9 and ICD10 coding: ICD9 410-414 or ICD10 I20-I25). This results in a holdout sample of *N* = 9,611 who meet the inclusion criteria and have PGIs for CVD. *N* = 4,061 (42.3%) of these individuals score in the top quintile of at least one CVD PGI.

## 1.6. G×E

Supplementary Tables 3 and 4 show the results of a regression of the G×E model explaining educational attainment. The independent variables are the respective PGIs, year of birth and their interaction. In Supplementary Table 3, the PGIs are used as continuous variables and in Supplementary Table 4 they are split into quintiles (using binary indicators). Supplementary Table 5 shows the results of pairwise *F*-tests that test whether the G×E coefficients are equal to each other.

|  | (1) | (2) | (3) | (4) | (5) | (6) |
| --- | --- | --- | --- | --- | --- | --- |
| Year of birth (YoB) | 0.15*** | 0.14*** | 0.14*** | 0.14*** | 0.14*** | 0.14*** |
|  | (0.00) | (0.00) | (0.00) | (0.00) | (0.00) | (0.00) |
| EA PGI 23andMe + UKB (LDpred) | 34.42*** |  |  |  |  |  |
|  | (6.35) |  |  |  |  |  |
| YoB × EA PGI 23andMe + UKB (LDpred) | -0.02*** |  |  |  |  |  |
|  | (0.00) |  |  |  |  |  |
| EA PGI 23andMe (LDpred) |  | 18.65*** |  |  |  |  |
|  |  | (6.56) |  |  |  |  |
| YoB × EA PGI 23andMe (LDpred) |  | -0.01*** |  |  |  |  |
|  |  | (0.00) |  |  |  |  |
| EA PGI UKB (LDpred) |  |  | 45.53*** |  |  |  |
|  |  |  | (6.41) |  |  |  |
| YoB × EA PGI UKB (LDpred) |  |  | -0.02*** |  |  |  |
|  |  |  | (0.00) |  |  |  |
| EA PGI 23andMe + UKB (C+T) |  |  |  | 32.86*** |  |  |
|  |  |  |  | (6.48) |  |  |
| YoB × EA PGI 23andMe + UKB (C+T) |  |  |  | -0.02*** |  |  |
|  |  |  |  | (0.00) |  |  |
| EA PGI 23andMe (C+T) |  |  |  |  | 13.14** |  |
|  |  |  |  |  | (6.62) |  |
| YoB × EA PGI 23andMe (C+T) |  |  |  |  | -0.01* |  |
|  |  |  |  |  | (0.00) |  |
| EA PGI 23andMe (C+T) |  |  |  |  |  | 44.31*** |
|  |  |  |  |  |  | (6.53) |
| YoB × EA PGI 23andMe (C+T) |  |  |  |  |  | -0.02*** |
|  |  |  |  |  |  | (0.00) |
| Observations | 38,049 | 38,049 | 38,049 | 38,049 | 38,049 | 38,049 |
| R-squared | 0.11 | 0.09 | 0.11 | 0.10 | 0.08 | 0.10 |

**Supplementary Table 3.** Results of the regressions explaining educational attainment (EA) for the sample of siblings of the UK Biobank. Robust standard errors in parentheses. All regressions control for the first 40 principal components of the genetic relatedness matrix. *** *p*<0.01, ** *p*<0.05, * *p*<0.10.

|  | (1) | (2) | (3) | (4) | (5) | (6) |
| --- | --- | --- | --- | --- | --- | --- |
| Year of birth (YoB) | 0.14*** | 0.15*** | 0.14*** | 0.16*** | 0.15*** | 0.14*** |
|  | (0.01) | (0.01) | (0.01) | (0.01) | (0.01) | (0.01) |
| EA PGI 23andMe + UKB (LDPRED) = 1 | -33.52 |  |  |  |  |  |
|  | (21.76) |  |  |  |  |  |
| EA PGI 23andMe + UKB (LDPRED) = 2 | -37.86* |  |  |  |  |  |
|  | (22.02) |  |  |  |  |  |
| EA PGI 23andMe + UKB (LDPRED) = 4 | -1.35 |  |  |  |  |  |
|  | (21.69) |  |  |  |  |  |
| EA PGI 23andMe + UKB (LDPRED) = 5 | 57.89*** |  |  |  |  |  |
|  | (20.65) |  |  |  |  |  |
| EA PGI 23andMe + UKB (LDPRED) = 1 × YoB | 0.02 |  |  |  |  |  |
|  | (0.01) |  |  |  |  |  |
| EA PGI 23andMe + UKB (LDPRED) = 2 × YoB | 0.02* |  |  |  |  |  |
|  | (0.01) |  |  |  |  |  |
| EA PGI 23andMe + UKB (LDPRED) = 4 × YoB | 0.00 |  |  |  |  |  |
|  | (0.01) |  |  |  |  |  |
| EA PGI 23andMe + UKB (LDPRED) = 5 × YoB | -0.03*** |  |  |  |  |  |
|  | (0.01) |  |  |  |  |  |
| EA PGI 23andMe (LDPRED) = 1 |  | -13.71 |  |  |  |  |
|  |  | (21.81) |  |  |  |  |
| EA PGI 23andMe (LDPRED) = 2 |  | -2.05 |  |  |  |  |
|  |  | (22.02) |  |  |  |  |
| EA PGI 23andMe (LDPRED) = 4 |  | 2.44 |  |  |  |  |
|  |  | (21.76) |  |  |  |  |
| EA PGI 23andMe (LDPRED) = 5 |  | 50.91** |  |  |  |  |
|  |  | (20.96) |  |  |  |  |
| EA PGI 23andMe (LDPRED) = 1 × YoB |  | 0.01 |  |  |  |  |
|  |  | (0.01) |  |  |  |  |
| EA PGI 23andMe (LDPRED) = 2 × YoB |  | 0.00 |  |  |  |  |
|  |  | (0.01) |  |  |  |  |
| EA PGI 23andMe (LDPRED) = 4 × YoB |  | -0.00 |  |  |  |  |
|  |  | (0.01) |  |  |  |  |
| EA PGI 23andMe (LDPRED) = 5 × YoB |  | -0.03** |  |  |  |  |
|  |  | (0.01) |  |  |  |  |
| EA PGI UKB (LDPRED) = 1 |  |  | -57.49*** |  |  |  |
|  |  |  | (21.81) |  |  |  |
| EA PGI UKB (LDPRED) = 2 |  |  | -23.43 |  |  |  |
|  |  |  | (22.11) |  |  |  |
| EA PGI UKB (LDPRED) = 4 |  |  | 20.60 |  |  |  |
|  |  |  | (21.50) |  |  |  |
| EA PGI UKB (LDPRED) = 5 |  |  | 65.77*** |  |  |  |
|  |  |  | (20.93) |  |  |  |
| EA PGI UKB (LDPRED) = 1 × YoB |  |  | 0.03** |  |  |  |
|  |  |  | (0.01) |  |  |  |
| EA PGI UKB (LDPRED) = 2 × YoB |  |  | 0.01 |  |  |  |
|  |  |  | (0.01) |  |  |  |
| EA PGI UKB (LDPRED) = 4 × YoB |  |  | -0.01 |  |  |  |
|  |  |  | (0.01) |  |  |  |
| EA PGI UKB (LDPRED) = 5 × YoB |  |  | -0.03*** |  |  |  |
|  |  |  | (0.01) |  |  |  |
| EA PGI 23andMe + UKB (C+T) = 1 |  |  |  | 14.87 |  |  |
|  |  |  |  | (21.88) |  |  |
| EA PGI 23andMe + UKB (C+T) = 2 |  |  |  | -2.26 |  |  |
|  |  |  |  | (21.90) |  |  |
| EA PGI 23andMe + UKB (C+T) = 4 |  |  |  | 40.91* |  |  |
|  |  |  |  | (21.65) |  |  |
| EA PGI 23andMe + UKB (C+T) = 5 |  |  |  | 93.07*** |  |  |
|  |  |  |  | (20.87) |  |  |
| EA PGI 23andMe + UKB (C+T) =1 × YoB |  |  |  | -0.01 |  |  |
|  |  |  |  | (0.01) |  |  |
| EA PGI 23andMe + UKB (C+T) =2 × YoB |  |  |  | 0.00 |  |  |
|  |  |  |  | (0.01) |  |  |
| EA PGI 23andMe + UKB (C+T) =4 × YoB |  |  |  | -0.02* |  |  |
|  |  |  |  | (0.01) |  |  |
| EA PGI 23andMe + UKB (C+T) =5 × YoB |  |  |  | -0.05*** |  |  |
|  |  |  |  | (0.01) |  |  |
| EA PGI 23andMe (C+T) = 1 |  |  |  |  | 13.30 |  |
|  |  |  |  |  | (22.16) |  |
| EA PGI 23andMe (C+T) = 2 |  |  |  |  | 30.68 |  |
|  |  |  |  |  | (22.08) |  |
| EA PGI 23andMe (C+T) = 4 |  |  |  |  | 21.76 |  |
|  |  |  |  |  | (22.02) |  |
| EA PGI 23andMe (C+T) = 5 |  |  |  |  | 63.23*** |  |
|  |  |  |  |  | (21.48) |  |
| EA PGI 23andMe (C+T) = 1 × YoB |  |  |  |  | -0.01 |  |
|  |  |  |  |  | (0.01) |  |
| EA PGI 23andMe (C+T) = 2 × YoB |  |  |  |  | -0.02 |  |
|  |  |  |  |  | (0.01) |  |
| EA PGI 23andMe (C+T) = 4 × YoB |  |  |  |  | -0.01 |  |
|  |  |  |  |  | (0.01) |  |
| EA PGI 23andMe (C+T) = 5 × YoB |  |  |  |  | -0.03*** |  |
|  |  |  |  |  | (0.01) |  |
| EA PGI UKB (C+T) = 1 |  |  |  |  |  | -61.15*** |
|  |  |  |  |  |  | (21.94) |
| EA PGI UKB (C+T) = 2 |  |  |  |  |  | -50.49** |
|  |  |  |  |  |  | (22.05) |
| EA PGI UKB (C+T) = 4 |  |  |  |  |  | -6.75 |
|  |  |  |  |  |  | (21.83) |
| EA PGI UKB (C+T) = 5 |  |  |  |  |  | 52.21** |
|  |  |  |  |  |  | (21.17) |
| EA PGI UKB (C+T) = 1 × YoB |  |  |  |  |  | 0.03*** |
|  |  |  |  |  |  | (0.01) |
| EA PGI UKB (C+T) = 2 × YoB |  |  |  |  |  | 0.03** |
|  |  |  |  |  |  | (0.01) |
| EA PGI UKB (C+T) = 4 × YoB |  |  |  |  |  | 0.00 |
|  |  |  |  |  |  | (0.01) |
| EA PGI UKB (C+T) = 5 × YoB |  |  |  |  |  | -0.03** |
|  |  |  |  |  |  | (0.01) |
| Observations | 38,049 | 38,049 | 38,049 | 38,049 | 38,049 | 38,049 |
| *R*-squared | 0.11 | 0.09 | 0.10 | 0.10 | 0.08 | 0.09 |

**Supplementary Table 4.** Results of the regressions explaining educational attainment (EA) for the sample of siblings of the UK Biobank. Robust standard errors in parentheses. All regressions control for the first 40 principal components of the genetic relatedness matrix. *** *p*<0.01, ** *p*<0.05, * *p*<0.10.

|  | *F*-statistic | *p* value |
| --- | --- | --- |
| *Joint F-test* |  |  |
| YOB×EA PGI 23andMe+UKB (LDPRED)= YOB×EA PGI 23andMe (LDPRED) = YOB×EA PGI UKB (LDPRED) = YOB×EA PGI 23andMe+UKB (C+T) = YOB×EA PGI 23andMe (C+T) = YOB×EA PGI UKB (C+T) | 3.55 | 0.003 |
| *Pairwise F-test* |  |  |
| YOB×EA PGI 23andMe+UKB (LDPRED)= YOB×EA PGI 23andMe (LDPRED) | 0.023 | 0.880 |
| YOB×EA PGI 23andMe+UKB (LDPRED)= YOB×EA PGI UKB (LDPRED) | 2.524 | 0.112 |
| YOB×EA PGI 23andMe+UKB (LDPRED) = YOB×EA PGI 23andMe+UKB (C+T) | 4.546 | 0.033 |
| YOB×EA PGI 23andMe+UKB (LDPRED) = YOB×EA PGI 23andMe (C+T) | 1.311 | 0.252 |
| YOB×EA PGI 23andMe+UKB (LDPRED) = YOB×EA PGI UKB (C+T) | 1.053 | 0.305 |
| YOB×EA PGI 23andMe (LDPRED) = YOB×EA PGI UKB (LDPRED) | 2.056 | 0.152 |
| YOB×EA PGI 23andMe (LDPRED) = YOB×EA PGI 23andMe+UKB (C+T) | 3.905 | 0.048 |
| YOB×EA PGI 23andMe (LDPRED) = YOB×EA PGI 23andMe (C+T) | 1.670 | 0.196 |
| YOB×EA PGI 23andMe (LDPRED) = YOB×EA PGI UKB (C+T) | 1.377 | 0.241 |
| YOB×EA PGI UKB (LDPRED) = YOB×EA PGI 23andMe+UKB (C+T) | 0.297 | 0.586 |
| YOB×EA PGI UKB (LDPRED) = YOB×EA PGI 23andMe (C+T) | 7.415 | 0.006 |
| YOB×EA PGI UKB (LDPRED) = YOB×EA PGI UKB (C+T) | 6.766 | 0.009 |
| YOB×EA PGI 23andMe+UKB (C+T) = YOB×EA PGI 23andMe (C+T) | 10.636 | 0.001 |
| YOB×EA PGI 23andMe+UKB (C+T) = YOB×EA PGI UKB (C+T) | 9.851 | 0.002 |
| YOB×EA PGI 23andMe (C+T) = YOB×EA PGI UKB (C+T) | 0.013 | 0.908 |

**Supplementary Table 5.** *F*-test results of pairwise comparisons of the interaction coefficients reported in Supplementary Table 3.

#### 1.7. Simulations

In simulations, the environment can be controlled and so we can generate a “true” PGI. Our simulations intend to help answering three main questions: (i) what is the rank concordance between the true PGI and any given estimated PGI depending on the predictive power of this estimated PGI?; (ii) What is the predicted rank concordance between any two estimated PGIs depending on their predictive power?; and (iii) what is the rank concordance between the true PGI and an outcome for varying levels of SNP-based heritability?

In practice, the true (latent) PGI is not observed, and we work with approximations of the latent PGI that is measured with error. We assume that an estimated PGI is equal to the true latent PGI (denoted by ${PGI}^{*}$) plus some additive classical measurement error $\upsilon$:

$PGI={PGI}^{*}+\upsilon,$	(2)

with $\upsilon\sim N\left[ 0,\sigma_{\upsilon}^{2} \right]$. Based on the attenuation bias arising from measurement error in a linear regression of the outcome on the PGI, we can derive that^26^:

$\sigma_{\upsilon}^{2}=\sigma_{{PGI}^{*}}^{2}\left( \frac{\beta_{st}^{2}}{\hat{\beta_{st}^{2}}}-1 \right)$, (3)

where $\sigma_{{PGI}^{*}}^{2}$ is the variance of the true latent PGI, $\beta_{st}$is the true standardized coefficient, and $\hat{\beta_{st}}$is an estimated standardized coefficient from the literature. We derive the “true” $\beta_{st}$from the square root of the SNP-based heritability estimate, and we infer $\hat{\beta_{st}}$from the square root from an estimated incremental *R^2^* in the literature. For example, the incremental *R^2^* of the EA PGI in the latest large-scale GWAS of EA is around 12%^5^, and thus $\hat{\beta_{st}}=\sqrt{0.12}\approx0.35.$ Then, if we assume that $\sigma_{{PGI}^{*}}^{2}=1$, the implied variance of measurement error of this estimate is $\sigma_{\upsilon}^{2}={0.5}^{2}/{0.35}^{2}-1 \approx1.04.$

To model a realistic variance for the measurement error $\sigma_{\upsilon}^{2}$, we calibrate the simulations based on the predictive power of existing PGIs for EA, but our conclusions hold for other complex traits at varying levels of heritability. The SNP-based heritability serves as the natural upper bound for the *R^2^* of the PGI^27^, which is approximately equal to 25% for EA ^4,28,29^. This corresponds to a correlation coefficient of 0.5 between the PGI and the outcome (i.e., the square root of the SNP-based heritability). Hence, we assume that the standardized outcome EA and the standardized true latent ${PGI}^{*}$are drawn from a bivariate standard normal distribution with correlation 0.5. In the simulations we therefore set the true standardised effect size to $\beta_{st}=0.5$ and $\sigma_{{PGI}^{*}}^{2}=1$. These choices are however largely inconsequential, because the bias of any given PGI depends on the ratio

$\frac{{\hat{\beta_{st}}}^{2}}{{\beta_{st}}^{2}}=\frac{R_{PGI}^{2}}{h_{SNP}^{2}}$ , (4)

which we coin “*explained* SNP-based heritability”^26^. In other words, what matters for the bias in a given PGI is how closely the estimated $R^{2}$ on basis of a given PGI (i.e., the square of the estimated standardized effect in a univariate regression) approximates the SNP-based heritability.

We run a simulation trial with a sample size *N=*100,000 and then generate (i) an outcome with varying levels of SNP-based heritability from 0 to 100 under the assumption that we know the true latent ${PGI}^{*}$, and (ii) one or more PGIs with varying levels of measurement error where the explained SNP-based heritability varies between 0 and 100. In turn we compute deciles and quintiles of the outcome, the true PGI and all generated PGIs, and assess the rank concordance between them in various ways as evidenced in the results. The intention is to gauge how measurement error affects rank concordance, holding the discovery sample and construction method constant.

Supplementary Figure 1 shows the expected rank concordance for two PGIs with varying levels of explained SNP-based heritability. In line with our empirical results, for two PGIs that each explain 50% of the SNP-based heritability, there will be a substantial lack of concordance in the respective rankings. With increasing explained SNP-based heritability, this discordance gradually disappears. Hence, increases in GWAS discovery samples and methodological advances in PGI construction will improve the rank concordance of different PGIs.

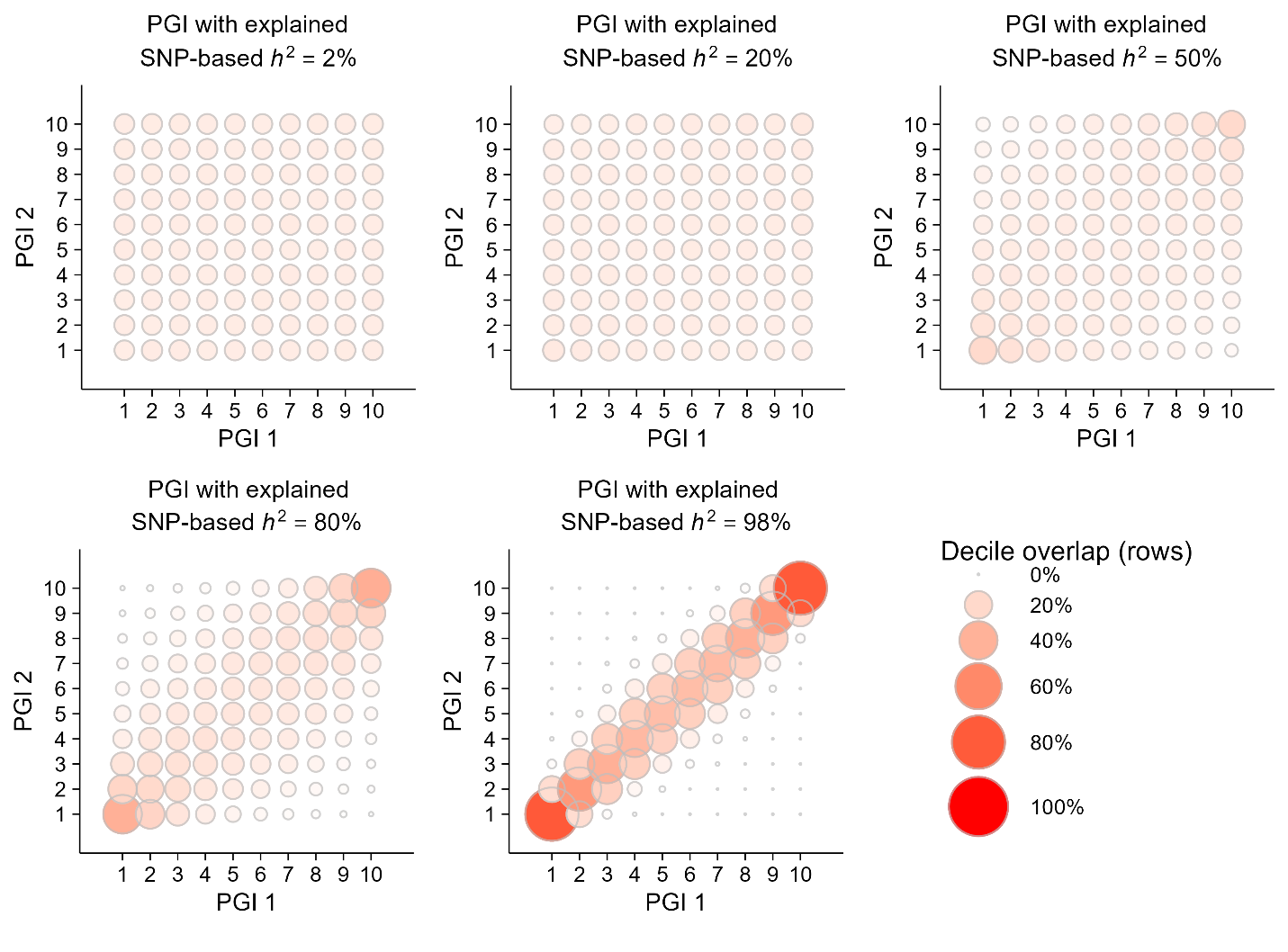

**Supplementary Figure 1.** The relationship between the rank in the distribution between two PGIs with the same level of explained SNP-based heritability (the $R^{2}$ of the PGI divided by the SNP-based heritability).

To quantify the on- and off-diagonal elements into one metric, and to show a more granular increase in explained SNP-based heritability, we now analyse the Spearman rank correlation between the estimated and “true” PGI. Supplementary Figure 2 shows the Spearman rank correlation between the “true” PGI and a PGI with varying levels of explained SNP-based heritability. The rank correlation is a concave increasing function of the explained SNP-based heritability.

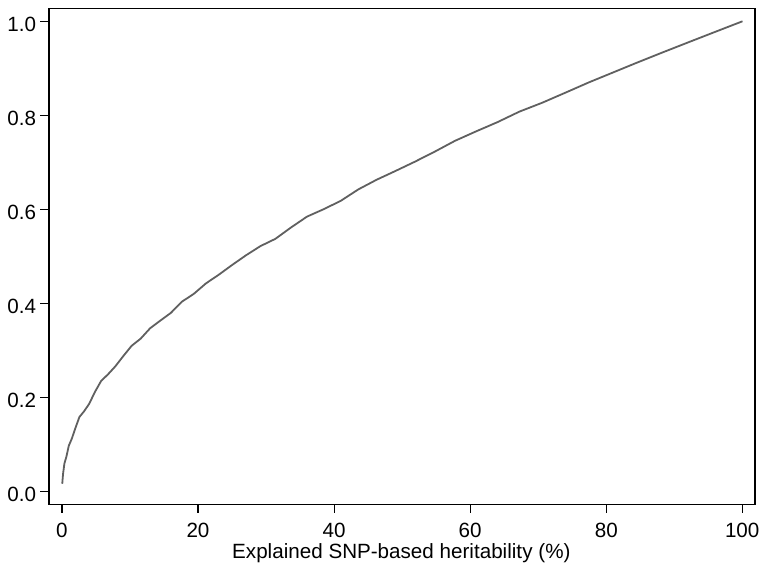

**Supplementary Figure 2.** The relationship between the Spearman correlation (rank of the “true” and estimated PGI) and the explained SNP-based heritability.

Finally, in Supplementary Figure 3 we show the relationship between the SNP-based heritability and the fraction correctly classified in the top quintile *of the trait distribution*. For a trait with a SNP-based heritability of 0, predicting the outcome based on a PGI would be a guess, with a random 20% classified correctly in the top quintile of the phenotype. What is more striking is how gradual this fraction rises as the SNP-based heritability increases. Even for traits with a SNP-based heritability of 75% (most traits will have heritability far below that), only about 60% of individuals is correctly classified in the top quintile on basis of genetic information alone. Hence, even when the “true” PGI would be observed for highly heritable traits, there will be a large fraction of individuals that are misclassified when it comes to predicting quintiles of the trait itself. Of course, when used in conjunction with environmental determinants of the trait, the true PGI can be a useful addition to the correct classification of risk within individuals. The below figure is therefore meant as a cautionary tale that there are two layers of uncertainty when using PGIs for risk prediction: first, any estimated PGI is a noisy proxy for the “true” PGI, and second, any risk prediction of even the “true” PGI is limited by the SNP-based heritability.

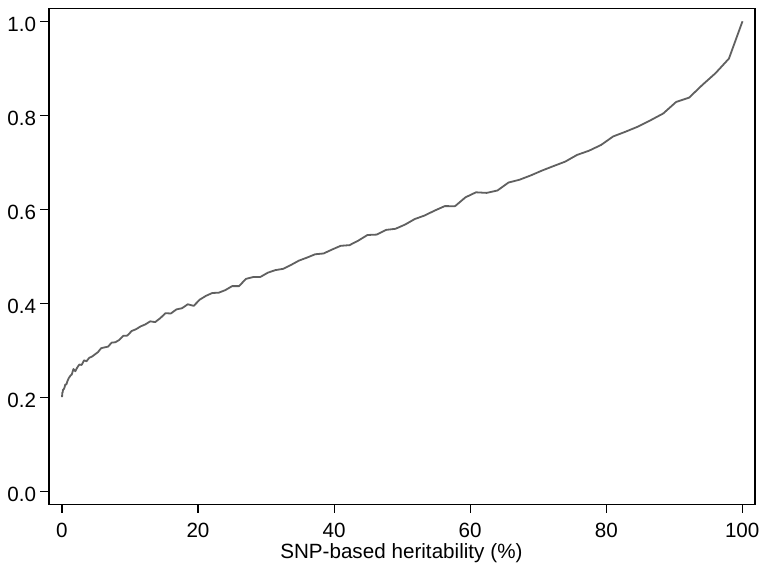

**Supplementary Figure 3.** The relationship between the SNP-based heritability (%) and the fraction of correctly classified individuals in the top quintile of the phenotype distribution when the “true” PGI is known.

### 2. References

1.	Fry, A. *et al.* Comparison of sociodemographic and health-related characteristics of UK biobank participants with those of the general population. *Am. J. Epidemiol.* **186**, 1026–1034 (2017).

2.	Bycroft, C. *et al.* The UK Biobank resource with deep phenotyping and genomic data. *Nature* **562**, 203–209 (2018).

3.	Manichaikul, A. *et al.* Robust relationship inference in genome-wide association studies. *Bioinformatics* **26**, 2867–2873 (2010).

4.	Okbay, A. *et al.* Genome-wide association study identifies 74 loci associated with educational attainment. *Nature* **533**, 539–542 (2016).

5.	Lee, J. J. *et al.* Gene discovery and polygenic prediction from a genome-wide association study of educational attainment in 1.1 million individuals. *Nat. Genet.* **50**, 1112–1121 (2018).

6.	Rietveld, C. A. *et al.* GWAS of 126,559 individuals identifies genetic variants associated with educational attainment. *Science* **340**, 1467–1471 (2013).

7.	Jiang, L. *et al.* A resource-efficient tool for mixed model association analysis of large-scale data. *Nat. Genet.* **51**, 1749–1755 (2019).

8.	Nikpay, M. *et al.* A comprehensive 1,000 genomes-based genome-wide association meta-analysis of coronary artery disease. *Nat. Genet.* **47**, 1121–1130 (2015).

9.	Winkler, T. W. *et al.* Quality control and conduct of genome-wide association meta-analyses. *Nat. Protoc.* **9**, 1192–1212 (2014).

10.	Willer, C. J., Li, Y. & Abecasis, G. R. METAL: fast and efficient meta-analysis of genomewide association scans. *Bioinformatics* **26**, 2190–2191 (2010).

11.	Benjamin, D., Cesarini, D., Laibson, D. I. & Turley, P. Social science genetics: a primer and progress report. *J. Econ. Lit.* (2020).

12.	Mills, M. C., Barban, N. & Tropf, F. C. An introduction to statistical genetic data analysis. *Cambridge MIT Press.* (2020).

13.	Becker, J. *et al.* Resource profile and user guide of the polygenic index repository. *Nat. Hum. Behav.* **5**, 1744–1758 (2021).

14.	Dudbridge, F. Power and predictive accuracy of polygenic risk scores. *PLoS Genet.* **9**, 1003348 (2013).

15.	DiPrete, T. A., Burik, C. A. P. & Koellinger, P. D. Genetic instrumental variable regression: explaining socioeconomic and health outcomes in nonexperimental data. *Proc. Natl. Acad. Sci. U. S. A.* **115**, 4970–4979 (2018).

16.	Smith-Woolley, E. *et al.* Differences in exam performance between pupils attending selective and non-selective schools mirror the genetic differences between them. *npj Sci. Learn.* **3**, 1–7 (2018).

17.	Davey Smith, G. & Ebrahim, S. ‘Mendelian Randomization’: Can Genetic Epidemiology Contribute to Understanding Environmental Determinants of Disease? *Int. J. Epidemiol.* **32**, 1–22 (2003).

18.	Privé, F., Vilhjálmsson, B. J., Aschard, H. & Blum, M. G. B. Making the most of clumping and thresholding for polygenic scores. *Am. J. Hum. Genet.* **105**, 1213–1221 (2019).

19.	Chang, C. C. *et al.* Second-generation PLINK: rising to the challenge of larger and richer datasets. *Gigascience* **4**, 1–16 (2015).

20.	Vilhjálmsson, B. J. *et al.* Modeling linkage disequilibrium increases accuracy of polygenic risk scores. *Am. J. Hum. Genet.* **97**, 576–592 (2015).

21.	Privé, F., Arbel, J. & Vilhjálmsson, B. J. LDpred2: better, faster, stronger. *Bioinformatics* **36**, 5424–5431 (2021).

22.	Purcell, S. *et al.* PLINK: a tool set for whole-genome association and population-based linkage analyses. *Am. J. Hum. Genet.* **81**, 559–575 (2007).

23.	Aragam, K. G. *et al.* Limitations of contemporary guidelines for managing patients at high genetic risk of coronary artery disease. *J. Am. Coll. Cardiol.* **75**, 2769–2780 (2020).

24.	Goff, D. C. *et al.* 2013 ACC/AHA guideline on the assessment of cardiovascular risk: a report of the american college of cardiology/american heart association task force on practice guidelines. *J. Am. Coll. Cardiol.* **63**, 2935–2959 (2014).

25.	Lloyd-Jones, D. M. *et al.* Prediction of lifetime risk for cardiovascular disease by risk factor burden at 50 years of age. *Circulation* **113**, 791–798 (2006).

26.	Van Kippersluis, H. *et al.* Using obviously-related instrumental variables to increase the predictive power of polygenic scores. *BioRxiv* (2020) doi:10.1101/2021.04.09.439157.

27.	Visscher, P. M., Hill, W. & Wray, N. R. Heritability in the Genomics Era — concepts and misconceptions. *Nat. Rev. Genet.* **9**, 255–266 (2008).

28.	Davies, G. *et al.* Genome-wide association study of cognitive functions and educational attainment in UK Biobank (N=112 151). *Mol. Psychiatry* **21**, 758–767 (2016).

29.	Tropf, F. C. *et al.* Hidden heritability due to heterogeneity across seven populations. *Nat. Hum. Behav.* **1**, 757–765 (2017).

30.	Kong, A. *et al.* The nature of nurture: effects of parental genotypes. *Science* **359**, 424–428 (2018).

31.	Plomin, R., DeFries, J. C. & Loehlin, J. C. Genotype-environment Interaction and Correlation in the Analysis of Human Behavior. *Psychol. Bull.* **84**, 309–322 (1977).

1. The limitations of polygenic indices are however also starting to become apparent. Perhaps most prominently, since the GWAS on basis of which the polygenic indices are constructed are very sparse on control variables, a polygenic index is not a clean measure of genetic endowments. Instead, an individual’s polygenic index will reflect the effects of his/her parental genotype, which could influence an individual’s outcome through environmental pathways – a phenomenon known as ‘genetic nurture’^30^ or ‘passive gene-environment correlation’^31^. Additionally, whereas the predictive power of polygenic indices is increasing and scores for many traits can now be meaningful to assess differences across groups of individuals, as we also highlight in this paper, predictions at the individual level are still virtually meaningless.
